# Adjuvanted subunit intranasal vaccine prevents SARS-CoV-2 onward transmission in hamsters

**DOI:** 10.1101/2024.05.13.593816

**Authors:** Yongjun Sui, Swagata Kar, Bhavna Chawla, Tanya Hoang, YuanKai Yu, Shannon M. Wallace, Hanne Andersen, Jay A. Berzofsky

**Author notes:** Corresponding authors: Yongjun Sui, Ph.D.

## Abstract

Most COVID-19 vaccine trials have focused on recipient protection, not protection of their contacts, a critical need. As a subunit intranasal COVID-19 vaccine reduced nasopharyngeal virus more than did an intramuscular (IM) vaccine, we hypothesized that this vaccine might reduce onward transmission to others. We vaccinated hamsters with either the IM-administrated Moderna mRNA vaccine twice or one dose of mRNA IM followed by adjuvanted subunit intranasal vaccine. 24 hours after SARS-CoV-2 challenge, these animals were housed with naïve recipients in a contactless chamber that allows airborne transmission. Onward airborne transmission was profoundly blocked: the donor and recipients of the intranasal vaccine-boosted group had lower oral and lung viral loads (VL), which correlated with mucosal ACE2 inhibition activity. These data strongly support the use of the intranasal vaccine as a boost to protect not only the vaccinated person, but also people exposed to the vaccinated person, a key public health goal.

**Author summary:** Natural transmission of SARS-CoV-2 is primarily airborne, through the respiratory mucosal route. However, current licensed COVID-19 vaccines are all intramuscular and induce more systemic than mucosal immunity. Here, we did a head-to-head comparison of COVID-19 booster vaccines on SARS-CoV-2 onward transmission. We found that compared to boosting with a Moderna mRNA systemic vaccine, a nanoparticle intranasal COVID-19 vaccine much more effectively prevents onward airborne transmission to naïve recipient hamsters. The protection was correlated with local mucosal antibody. Thus, a mucosal nanoparticle vaccine should be considered for preventing onward airborne transmission, a key public health necessity that has not been adequately studied.

## Introduction

Blocking viral transmission is an important function of efficient vaccines. From a public health point of view, preventing SARS-CoV-2 transmission to other susceptible individuals is extremely critical. However, most COVID-19 vaccine clinical trials studied only safety and protection of the vaccine recipient, but not prevention of transmission to others. Indeed, the currently licensed SARS-CoV-2 vaccines are successful to alleviate COVID-19-related hospitalization and deaths, but less effective against acquisition of infection and onward transmission. Though studies on SARS-CoV-2 breakthrough infections suggested that vaccine breakthrough infections are less contagious than primary infections in unvaccinated individuals[1, 2], the effects of these vaccines on reducing transmissibility have not been well evaluated.

As SARS-CoV-2 transmission is mostly through the nasopharynx, mucosal immunity could potentially reduce or abort the SARS-CoV-2 replication at the portal of entry (nasopharynx) to prevent virus from being transmitted to others. Intranasal administration of current vaccines, however, led to inconsistent results against SARS-CoV-2 infections [3, 4]. The adjuvant subunit mucosal vaccine, which induces vigorous mucosal immunity in the upper and lower respiratory tracts [5-7], and is more effective at clearing upper airway virus than a similar subunit vaccine given intramuscularly (IM), may have the potential to better reduce SARS-CoV-2 onward transmission. Here, we assessed whether the adjuvanted subunit vaccine delivered intranasally could protect from onward transmission of SARS-CoV-2 in a hamster model better than the systemic mRNA vaccine. As SARS-CoV-2 virus can be effectively transmitted among the hamsters, this represents a more natural dose and route of infection/transmission [8].

## Results and Discussion

To vaccinate the donor hamsters, we first primed IM two groups of male animals (n=5/group) with Moderna bivalent mRNA COVID-19 vaccine (Moderna Therapeutics, MA) to mimic the fact that many individuals have already been vaccinated with at least one or more doses of systemic vaccines. Male animals were chosen as they are more likely to have higher viral load (VL) and more severe disease (Fig.1A). Three weeks later, Group 1 was boosted with the same mRNA vaccine (IM), while Group 2 was boosted intranasally (IN) with CP-15 adjuvanted (CpG+polyI:C+IL15) spike protein in DOTAP. Four weeks later, all groups, including a naïve control group (Group 3), were intranasally challenged with SARS-CoV-2 virus.

**Figure 1.**
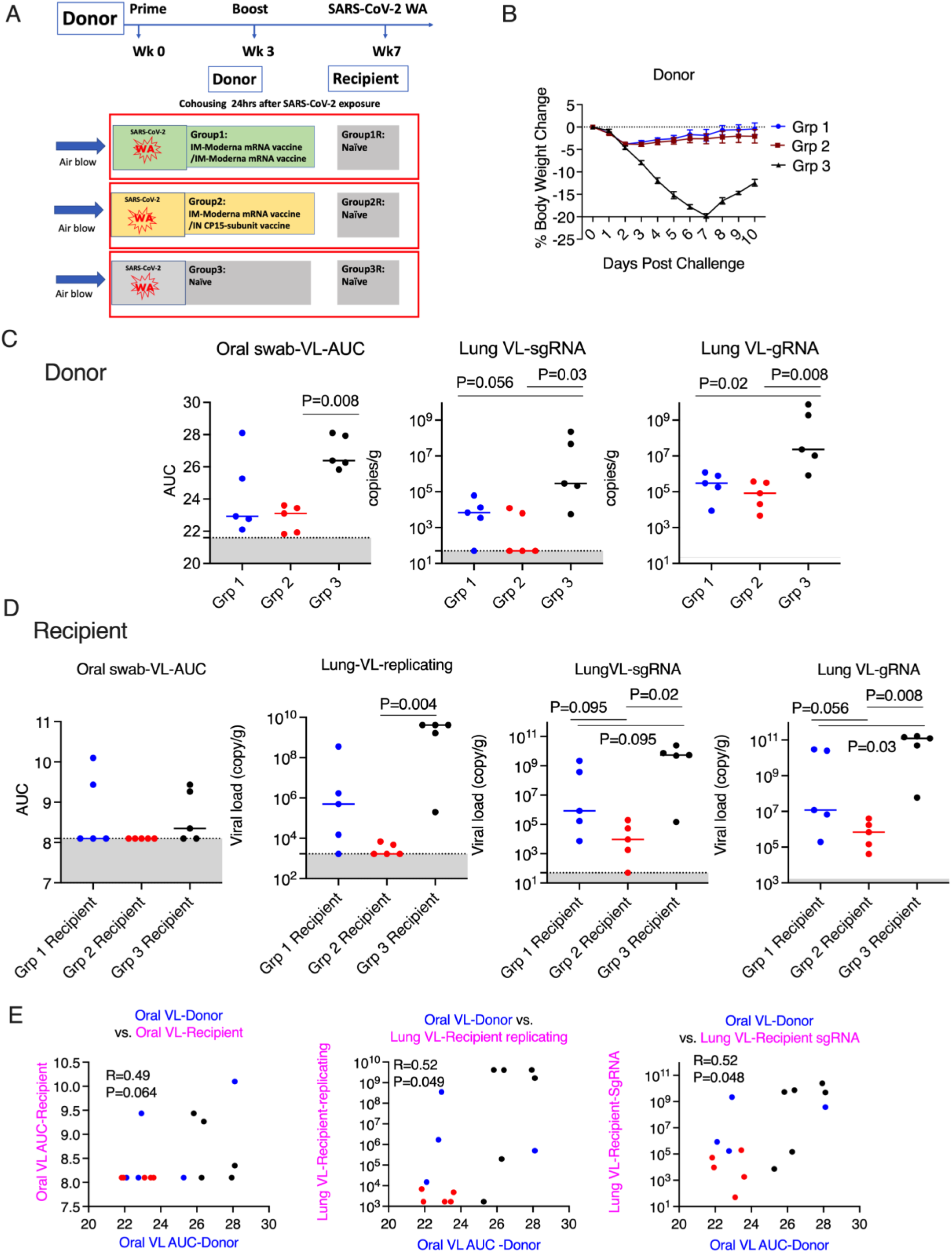
Mucosal vaccine prevented SARS-CoV-2 onward transmission in hamsters. A). Schematic of SARS-CoV-2 transmission study. B). Body weight change in the donor hamsters after the challenge of SARS-CoV-2 Washington strain. C). Area under curve (AUC) of viral load (VL) in oral swabs at Days 1, 2, 5, and 7, and sgRNA/ gRNA VL in the lung at Day 10 after viral challenge in the donor hamsters. D). AUC of VL in oral swabs at Day 1, 2, and 3, replicating, sgRNA/ gRNA VL in the lung at Day 3 after housing in the recipient hamsters. E). the Spearman’s correlations between the VL in the donor animals and the VL in the recipient animals. The groups are color coded, with Blue, red, and black denoting group 1-3 donor and recipient animals respectively.

Three groups of naïve hamsters (n=5/group) were used as recipients to assess the transmission rate 24 hours after the SARS-CoV-2 viral challenge. In each contact-free cage, one donor animal was housed with one naive recipient animal for 8 hours with unidirectional air flow, through a permeable membrane that prevented physical touching or transfer of secretions, but allowed airborne transmission of virus, from the donor to the recipient. The donor animals were monitored for weight loss, oral VL for an additional 8 days after housing, while the recipient groups were necropsied at day 3 post viral exposure to examine the VL in the lung tissues (Fig. 1A). Neither donor vaccine group showed significant weight loss, suggesting both vaccines as a booster could provide sufficient protection against disease (Fig. 1B). Both vaccine groups reduced viral replication (P=0.056, 0.03 for Group 1, 2) in the lung tissues (at day 10) compared to the naïve group. However, Group 2, but not Group 1, showed significant viral reduction in the oral swabs (Fig. 1C). For recipient groups, none housing with Group 2 had detectable oral VLs, indicating complete protection, while all 3 animals housing with Group 3 and 2 housing with Group 1 showed oral VLs (Fig. 1C). In the lung, the replicating virus titer and sgRNA were significantly reduced in the animals housing with Group 2 (P=0.004, P=0.02), while no significant no protection was observed in the animals housing with Group 1 (Fig. 1D). Nevertheless, both systemic and mucosal vaccines demonstrated significant reduction of SARS-CoV-2 gRNA, though the hamsters showed trend of lower gRNA level in the recipient group housing with group 2 than those with group 1 (Fig. 1C-D). These data indicated that as a booster, the mucosal vaccine provided substantially better blockage against onward transmission than the systemic mRNA vaccine did. Histopathological exams revealed that both vaccines were effective in reducing SARS-CoV-2-related microscopic findings in the lung when compared to Group 3 animals; Group 2 was most effective with the lowest incidence of findings (Table 1). Additionally, both vaccines were effective in eliminating the transmission of SARS-CoV-2 related microscopic findings in the lung of untreated co-housed animals (Table 1). We also found that oral VL in the donors was correlated with lung and oral VLs in the recipients (Fig. 1E), which was consistent with the previous finding that onward viral transmission is multifactorial, and the level of infectious virus was one of the key parameters [9].

**Table 1.** SARS-CoV-2-related microscopic findings in the lung of the donor (necropsied at Day 10 post challenge) and the recipient animals (necropsied at Day 3 post housing) *.

| **Groups** | **1** | **2** | **3** | **Groups** | **1 Recipient** | **2 Recipient** | **3 Recipient** |
| --- | --- | --- | --- | --- | --- | --- | --- |
| Animals/Group | 5 | 5 | 5<sup>a</sup> | Animals/Group | 5 | 5 | 5 |
| **Inflammation, mixed or mononuclear cell, alveolar or bronchoalveolar** | 3 | - | 5 | **Inflammation, mononuclear cell, alveolar** | - | - | 1 |
| minimal | 3 | - | - | minimal | - | - | 1 |
| mild | - | - | 4 |  |  |  |  |
| moderate | - | - | - |  |  |  |  |
| marked | - | - | 1 |  |  |  |  |
| **Inflammation, mononuclear cell, vascular / perivascular** | - | - | 5 | **Inflammation, mononuclear cell, vascular / perivascular** | - | - | 3 |
| minimal | - | - | 3 | minimal | - | - | 2 |
| mild | - | - | 2 | mild | - | - | 1 |
| **Fibrosis or fibroplasia, pleural** | - | - | 2 |  |  |  |  |
| minimal | - | - | 1 |  |  |  |  |
| mild | - | - | 1 |  |  |  |  |
| **Syncytial cell** | - | - | 5 |  |  |  |  |
| minimal | - | - | 5 |  |  |  |  |
| **Hyperplasia, bronchiolo-alveolar or alveolar** | 5 | 1 | 5 |  |  |  |  |
| minimal | 3 | - | - |  |  |  |  |
| mild | 2 | 1 | - |  |  |  |  |
| moderate | - | - | 3 |  |  |  |  |
| marked | - | - | 2 |  |  |  |  |
| **Hyperplasia, endothelial** | - | - | 2 |  |  |  |  |
| minimal | - | - | 2 |  |  |  |  |
| Hemorrhage | - | - | 2 |  |  |  |  |
| minimal | - | - | - |  |  |  |  |
| mild | - | - | 2 |  |  |  |  |
| **Hypertroplasia, mesothelial cell** | - | - | 1 |  |  |  |  |
| minimal | - | - | - |  |  |  |  |
| mild | - | - | 1 |  |  |  |  |

\*Findings were graded 1-5, depending upon severity. Microscopic findings, if applicable, are correlated with macroscopic observations. For severity grades, equivalent numbered grades are 1 = minimal, 2 = mild, 3 = moderate, 4 = marked, 5 = severe.

To delineate the mechanisms, we assessed the ACE2 inhibition activity (a surrogate neutralizing antibody assay) against the Wuhan and 9 Omicron sub-strains in the serum and oral swabs of the vaccinated animals. In the serum, Group 1 had similar or higher levels of ACE2 inhibition activity compared to Group 2 (Figure 2). However, in the oral swabs (Fig 3A), we found the opposite: Group 2 had significantly (or trending) higher ACE2 inhibition activity than Group 1 did for wild type, BA.1, BA.2.75, BA.5, BN.1, BQ.1, BQ.1.1, and XBB.1.5. Moreover, ACE2 inhibition activity in oral swabs was inversely correlated with oral VLs, suggesting the mucosal immunity might play a more important role in reducing viral infections (Fig. 3B). These data might also explain the observation that asymptomatic or mildly symptomatic breakthrough infections occurred in subjects who maintained their systemic antibody and T-cell responses [10].

**Figure. 2.**
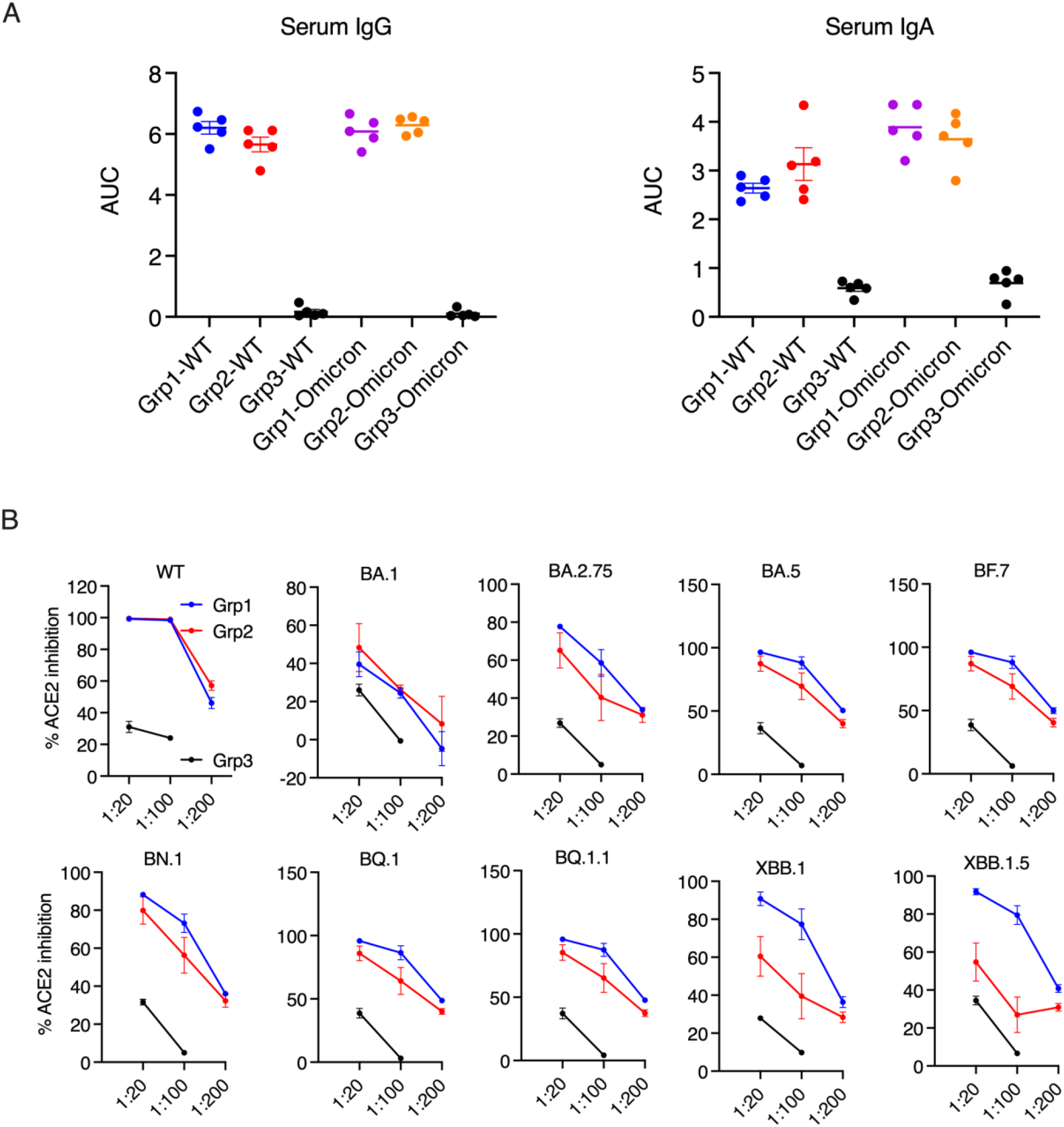
Serum antibody responses and ACE2 inhibition activity in the donor animals 2 weeks after the boost. A). Anti-spike IgG and IgA in serum samples were measured using ELISA. The serum samples were diluted from 1:100, 4-fold dilution, and 6 dilutions. Area under curve (AUC) of serum IgG and IgA are shown. B). The ACE2 inhibition activity against wild type (WT) and Omicron sub-strains in the serum samples of the donor animals (2 weeks after the boost).

**Figure 3.**
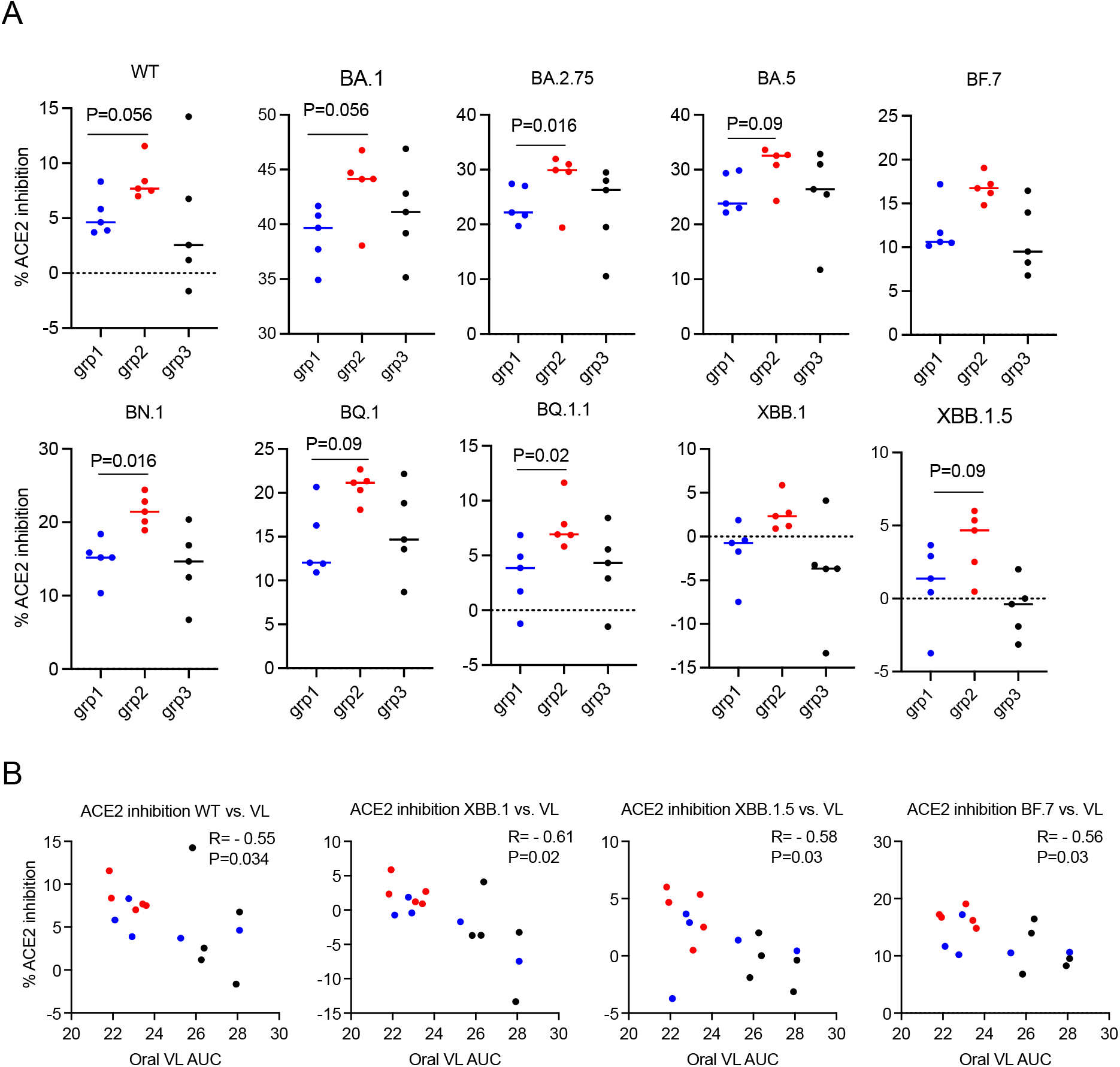
ACE2 inhibition activity was inversely correlated with viral load (VL) in the oral swabs. A). the ACE2 inhibition activity against wild type (WT) and Omicron sub-strains in the oral swabs of the donor animals (2 weeks after the boost). B). Spearman’s correlations between ACE2 inhibition activity against WT/Omicron sub-strains XBB. 1/XBB. 1.5/BF.7 and the viral load (VL) in oral swabs. The groups are color coded, with Blue, red, and black denoting group 1-3 donor animals respectively.

As most current vaccines prevent COVID-19 disease but do not prevent initial infection and spread to others, it is important to focus on developing vaccines that block SARS-CoV-2 transmission. Previous studies using adenovirus type 5 SARS-CoV-2 mucosal vaccines showed reducing viral transmission compared to naïve donors [11]. However, the data to compare the protection capacity with currently licensed systemic vaccines are lacking. Here we did a head-to-head comparison of a subunit mucosal vaccine versus a licensed mRNA vaccine for their ability to induce mucosal immunity and subsequently prevent airborne SARS-CoV-2 onward transmission from vaccinated donors to naïve recipients. Our results demonstrated that CP15 adjuvanted subunit vaccine, as a booster, mediated better protection against onward airborne transmission, compared to the licensed systemic mRNA SARS-CoV-2 vaccine, and the induction of mucosal naso-oropharyngeal immunity was a correlate of protection (Fig. 3B). Thus, this mucosal vaccine, along with other mucosal vaccines, could act as attractive vaccine booster candidates to limit SARS-CoV-2 onward transmission and fulfill a critical public health need that has not been addressed.

## Material and Methods

### Animals

All animal studies were approved by the Institutional Animal Care and Use Committee of BIOQUAL, Inc. (Rockville, MD. Thirty male Syrian golden hamsters (Envigo), 8–10 weeks old, were housed and conducted in compliance with all relevant regulations.

### Vaccination

The hamsters were grouped randomly into 6 groups (N=5/group). Group 1-3 were the donor groups, and Group 4-6 were the recipient groups. 10 µg/dose Moderna COVID-19 bivalent vaccine (in 100µl) was given to Group 1 -2 intramuscularly at Day 0. On Day21, Group1 got the same dose/route of Moderna COVID-19 vaccine. Group 2 received intranasally CP-15 adjuvanted mucosal vaccine, which was composed of 20 µg of SARS-CoV-2 Spike S1+S2 trimer protein (10 µg D614G + 10 µg B.1.1.529) (40589-V08H8, 40589-V08H26, Sino Biological. Inc.), mixed with 20 µg of D-type CpG oligodeoxynucleotide (vac-1826-1, In vivoGen), 40 µg of Poly I:C (vac-pic, In vivoGen), and 20 µg of recombinant murine IL-15 (210-15, PeproTech) in 20 µl of DOTAP (11 811 177 001, Roche Inc.). For the intranasal procedures, the hamsters were sedated with Ketamine (80µg/kg)/Xylazine(5µg/kg), and 50 uL/nare, total 100uL vaccine was administrated per hamster. The dosing material was more likely to penetrate further into the respiratory tract, getting to the lungs by using this procedure (sedation and size of the inoculum).

### Viral challenge and viral transmission

Four weeks after the last vaccination, Group 1-3 (Group 3 was the naïve control group) were challenged with 6×10^3^ PFU SARS-CoV-2 WAS-CALU-3 (LOT: 12152020-1235, BEI Resources). The animals were sedated, and virus challenge was given intranasally with 50 µl/nare as described before [7]. Body weights were monitored before and after the viral challenge. 24hrs after the viral challenge, each animal from Group1-3 was housed with one naïve hamster from Group 4-6 respectively. The housing was in a single contract-free transmission chamber as described before [11]. The chamber was designed so that the airflow was unidirectional: from the infected donor animal to the naïve recipient one. Animals were housed 1:1 with a transmission divider for 8 hours, then single housed. The donor animals were monitored for an additional 8 days for body weights, oral VLs. At Day10, the donor animals were necropsied, and lung viral loads were measured. For recipient groups, body weights and oral VLs was monitored and at Day3, the animals were necropsied, and lung VLs were measured.

### Viral load measurements

TCID50 assays were used to measure live virus as described before [7]. Briefly, 20 µL of sample was 10-fold serially diluted and added to Vero TMPRSS2 cells, cultured in DMEM + 2% FBS + Gentamicin at 37°C, 5.0% CO2 for 4 days. Virus stock of known infectious titer was included in the assay as a positive control, while medium only served as a negative control. Cytopathic effect (CPE) was inspected, and the TCID50 value was calculated using the Read-Muench formula. SARS-CoV-2 RNA levels were assessed by reverse transcription PCR at BIOQUAL Inc. as previously described [5]. RNA was extracted from oral swab and homogenized lung tissue samples. Subgenomic/viral RNA using different primer/probe sets, targeting the viral E gene mRNA or the viral nucleocapsid respectively, was measured. VLs are shown as copies per swab for oral samples, and copies per gram for lung tissues, with a cutoff value of 50 copies.

### ELISA and ACE2 inhibition assay

ELISA was performed as described [7]. The V-PLEX-SARS-CoV-2 Panel 32 (ACE2) Kit was used according to the manufacturer’s instructions using a Sector Imager 2400 (Meso Scale discovery). Serum samples (diluted 20-, 100-, and 200-fold), or oral samples (2-fold dilution) were added into the pre-coated V-plex plates. ACE2 binding and detection reagents were sequentially added.

### Histopathological exams

Groups 1-3 were euthanized on Day 10 post viral challenge. Recipient groups were euthanized on Day 3 post co-housing. At necropsy, the left lung was collected and placed in 10% neutral buffered formalin for histopathologic analysis. Tissue sections were trimmed and processed to hematoxylin and eosin (H&E) stained slides, all slides were examined by a board-certified pathologist and recorded using the Pristima® 7.5.0 Build 8 version computer system.

### Statistical analysis

Statistical analyses were performed using Prism version 9. Oral swab viral load was presented as area under curve (AUC) values. Mann-Whitney and Spearman analyses were used for group comparisons and correlations. All statistical tests were 2-tailed.

## Acknowledgments

This work was supported by funding Z01 BC-011941 to Jay A. Berzofsky from Intramural Research Program, Center for Cancer Research, National Cancer Institute, National Institutes of Health.

